# The giant sequoia genome and proliferation of disease resistance genes

**DOI:** 10.1101/2020.03.17.995944

**Authors:** Alison D. Scott, Aleksey V. Zimin, Daniela Puiu, Rachael Workman, Monica Britton, Sumaira Zaman, Madison Caballero, Andrew C. Read, Adam J. Bogdanove, Emily Burns, Jill Wegrzyn, Winston Timp, Steven L. Salzberg, David B. Neale

## Abstract

The giant sequoia (*Sequoiadendron giganteum*) of California are massive, long-lived trees that grow along the U.S. Sierra Nevada mountains. As they grow primarily in isolated groves within a narrow range, conservation of existing trees has been a national goal for over 150 years. Genomic data are limited in giant sequoia, and the assembly and annotation of the first giant sequoia genome has been an important goal to allow marker development for restoration and management. Using Illumina and Oxford Nanopore sequencing combined with Dovetail chromosome conformation capture libraries, 8.125 Gbp of sequence was assembled into eleven chromosome-scale scaffolds. This giant sequoia assembly represents the first genome sequenced in the Cupressaceae family, and lays a foundation for using genomic tools to aid in giant sequoia conservation and management. Beyond conservation and management applications, the giant sequoia assembly is a resource for answering questions about the life history of this enigmatic and robust species. Here we provide an example by taking an inventory of the large and complex family of NLR type disease resistance genes.

## INTRODUCTION

Giant sequoia, *Sequoiadendron giganteum* (Lindl.) J.Buchh., is a California endemic conifer found in fragmented groves throughout the U.S. Sierra Nevada mountain range. Giant sequoias are known for their substantial size; individual specimens can reach over 90 m in height, more than 10 m in diameter, and may exceed 1000 m^3^ of wood volume (Sillett et al., 2015). In addition to their considerable proportions, giant sequoias are among the oldest tree species, as individuals can live for over 3,200 years (Douglass, 1919). Giant sequoias are one of the two redwood species in California, where they share the title of state tree with their closest relative, the coast redwood (*Sequoia sempervirens* Endl.).

Though they have occupied their current range for millennia and were known by indigenous people for centuries before colonizers arrived, giant sequoias became icons of the American west beginning with the exploitation of the Discovery Tree in 1853 (Cook, 1961). Despite the brittle nature of their wood, historical research indicates a third of groves were either completely or partially logged (Elliot-Fisk et al., 1997, cited by Burns et al., 2018). Giant sequoias were first protected in 1864 (Cook, 1961), and have remained a cornerstone of the American conservation movement ever since.

While the majority (98%) of remaining giant sequoia groves are now protected (Burns et al., 2018), the species is listed as endangered (IUCN) and is overall experiencing a decline (Schmid & Farjon, 2013). The dwindling numbers of giant sequoia are largely attributed to a lack of reproductive success due in part to fire suppression over the last century (Stephenson, 1994), as giant sequoia trees rely on extreme heat to open their cones and release seeds in addition to preparing the understory for germination. Mature giant sequoias in natural stands appear to withstand most pests and diseases, but relatively little is documented about the potential impact of insects and pathogens on younger trees. Recent research suggests giant sequoias are potentially susceptible to bark beetles, which can exacerbate the impacts of drought (Stephenson et al., 2018).

In plants, disease resistance is typically conferred by genes encoding nucleotide binding leucine-rich repeat (NLR) proteins that individually mediate responses to different pathogens. In crop species, NLR genes have demonstrated contributions to resistance against insects (Stahl et al., 2018), and a recent examination of transcriptome data from several conifer species showed that many conifer NLRs were induced following drought stress (Van Ghelder et al., 2019), suggesting an even broader role. Their importance in resilience to disease and abiotic stress makes cataloging NLR genes of particular interest for conservation and management. Notably, however, across species and even among plant populations, NLR genes account for the majority of copy-number and presence/absence polymorphisms (Yu et al., 2011; Zheng et al., 2011; Xu et al., 2012; Bush et al., 2013; Schatz et al., 2014), and this complexity makes accurate inventory challenging in the absence of a high quality genome assembly.

More broadly, a whole genome reference assembly provides a foundation for understanding the distribution of genetic variation in a species, which is critical for conservation and management. Though studies of population genetics and phylogenetics of giant sequoia have been conducted using isozymes, microsatellites, RADseq, and transcriptomic data (Fins and Libby, 1982; DeSilva & Dodd, 2014; Dodd & DeSilva, 2016; Scott et al., 2016) there is a dearth of robust genomic resources in this species. The closest species with fully sequenced genomes exist entirely in the family Pinaceae, which last shared a common ancestor with giant sequoia (Cupressaceae) more than 300 million years ago (Leslie et al., 2018).

A combination of short-read Illumina data, long-read Oxford Nanopore data, and Dovetail proximity ligation libraries produced a highly contiguous assembly with chromosome-scale scaffolds, many of which are telomere-to-telomere. This assembly also includes the largest scaffolds assembled to date in any organism. The genome was found to contain over 900 complete or partial NLR genes, of which over 250 are in consensus with annotation derived from protein evidence and gene modeling. The giant sequoia genome assembly and annotation presented here is an unprecedented resource in conifer genomics, both for the quality of the assembly and because it represents an understudied branch of the gymnosperm tree of life.

## MATERIALS AND METHODS

### Sequencing and assembly

#### Megagametophyte DNA extraction and sequencing

Cones were collected from a 1,360-year-old giant sequoia (SEGI21, Sillett et al., 2015) in Sequoia/Kings Canyon National Park in 2012. As in previous conifer genome sequencing projects (e.g. Zimin et al., 2014), the megagametophyte from a single fertilized seed was dissected out and its haploid DNA extracted with a Qiagen DNeasy Plant Kit (Hilden, Germany), followed by library preparation with an Illumina TruSeq Nano kit using the low throughput protocol. This megagametophyte library was then sequenced on 10 lanes of an Illumina HiSeq 4000 with 150 bp paired-end reads at the UC Davis Genome Center DNA Technologies Core facility.

#### Foliage DNA extraction and Nanopore sequencing

In 2017 foliage was collected from the upper canopy of the same giant sequoia tree (SEGI21). From this foliage, high molecular weight DNA was extracted following the protocol described here (dx.doi.org/10.17504/protocols.io.4vbgw2n). Briefly, purified genomic DNA was isolated through a nuclei extraction and lysis protocol. First, mature leaf tissue was homogenized in liquid nitrogen until well-ground, then added to a gentle lysis buffer (after Zhang et al., 2016, containing spermine, spermidine, triton, and β-mercaptoethanol) and stirred at 4°C for ten minutes. Cellular homogenate was filtered through five layers of Miracloth into a 50mL Falcon tube, then centrifuged at 4°C for 20 minutes at 1900 x g, which was selected based on the estimated giant sequoia genome size of around 9 Gb (Zhang et al., 2012; Hizume et al., 2001). Extracted nuclei were then lysed and gDNA precipitated using the Circulomics Nanobind Plant Nuclei Big DNA kit - alpha version (SKU NB-900-801-01). Then 1 μg of purified genomic DNA was input into the Ligation sequencing kit (LSK108-LSK109, Oxford Nanopore), according to protocol, with the exception of end repair optimization (100 μL sample, 14 μL enzyme, 6 μL enzyme at 20°C for 20 minutes, then 65°C for 20 minutes). Samples were sequenced on R9.4 minION flowcells using either the minION or GridION for 48 hours, then raw fast5 data was basecalled with Albacore version 2.13.

#### Hi-C and Chicago library preparation and sequencing

Additional foliage from SEGI21 was submitted to Dovetail Genomics (Scotts Valley, CA) for Hi-C and Chicago library preparation as described by Putnam et al., 2016. Hi-C libraries preserve *in vivo* chromatin structures while Chicago libraries are based on *in vitro* reconstituted chromatin; the combination of these two approaches allows for marked improvement in contiguity for genome assemblies. Three Hi-C libraries and two Chicago libraries passed QC for sequencing and were sent to the UC San Francisco Center for Advanced Technology where they were pooled and sequenced on an Illumina Novaseq 6000 in a single lane of an S4 flowcell (PE 150 bp).

#### Genome assembly

Assembly of the giant sequoia genome involved two major steps: contig assembly from Illumina and Oxford Nanopore reads and scaffolding with Chicago and Hi-C data by Dovetail Genomics. Contigs were produced using MaSuRCA assembler version 3.2.4 (Zimin et al, 2013, Zimin et al, 2017) with the default parameters. Then the sequence data from the two Chicago libraries were used to scaffold the initial contig assembly using Dovetail’s HiRise software (Putnam et al., 2016). Following this step, the output assembly comprised of Illumina, Oxford Nanopore, and Chicago data plus the Hi-C data was used as input for a second run of HiRise re-scaffolding software. The initial contig assembly was named giant sequoia 1.0 and the final scaffolded assembly giant sequoia 2.0.

#### Identification of centromeric and telomeric repeats

Tandem repeat elements up to 500 bp long were identified with the tandem repeat finder program (trf v4.09; Benson, 1999) with the recommended parameters (max mismatch delta PM PI minscore maxperiod, 2 7 7 80 10 50 500 resp.). A histogram of repeat unit lengths was then produced, which had the peaks at 7, 181, and 359 bp.

### Annotation

#### RNA isolation and sequencing

RNA was isolated from giant sequoia roots, foliage, and cambium using a LiCl-Urea buffer followed by cleanup using Zymo columns and reagents (Zymo Research, Irvine, CA). RNA quality was assessed using an Experion Electrophoresis System (Bio-Rad, Hercules, CA) and Qubit fluorometer (Thermo Fisher Scientific, Waltham, MA).

Double-stranded cDNA was generated from total RNA (2 µg per tissue) using the Lexogen TeloTM prime Full-length cDNA Kit (Lexogen, Inc., Greenland, NH, USA). Tissue-specific cDNAs were first barcoded by PCR (16-19 cycles) using IDT barcoded primers (Integrated DNA Technologies, Inc., Coralville, Iowa), and then bead-size selected with AMPure PB beads (two different size fractions of 1X and 0.4X). The three cDNAs were pooled in equimolar ratios and used to prepare a SMRTbell™ library using the PacBio Template Prep Kit (PacBio, Menlo Park, CA). The SMRTbell™ library was then sequenced on a Sequel v2 SMRT cell with polymerase 2.1 and chemistry 2.1 (P2.1C2.1) on one PacBio Sequel v2 SMRT cell at the UC Davis Genome Center DNA Technologies Core Facility.

#### Processing of IsoSeq data

Raw IsoSeq subreads were processed using the PacBio IsoSeq3 v3.0 workflow (https://github.com/PacificBiosciences/IsoSeq/blob/master/README_v3.0.md). Briefly, ccs v.3.0.0 was run to merge subreads one full-length circular consensus sequence (ccs) per Zero Mode Waveguide (ZMW). Then, lima v.1.7.0 was run to remove primer artifacts and to demultiplex the ccs by library barcode. Finally, isoseq3 cluster 3.0.0 was run to cluster the demultiplexed CCS reads into transcripts.

#### Repetitive element library generation and masking

RepeatModeler (2.0; Smit and Hubley, 2008) was used to detect *de novo* repeats in the giant sequoia 2.0 assembly, after scaffolds shorter than 3 kbp were removed. The resulting repeat library with classification was used as input for RepeatMasker (v4.0.9, Smit, Hubley, and Green, 2013) which soft masks repetitive elements in the genome. After this initial repeat masking using the de novo giant sequoia repeat library, RepeatMasker was run using a library of conifer repeats identified in other gymnosperm species clustered at 80% to further mask repetitive elements.

#### Structural annotation

PacBio IsoSeq data and previously published Illumina RNAseq data (Scott et al., 2016) were mapped to the soft masked genome, using Minimap2 v.2.12 (Li, 2018) for the long-read data and HISAT2 v.2.1.0 (Kim, Langmead, and Salzberg, 2015) for short reads. The resulting alignment files were merged and sorted, then used alongside protein evidence generated with GenomeThreader (Gremme et al., 2005) as input to Braker2 v2.1.2 (Hoff et al., 2019; Hoff et al., 2015; Stanke et al., 2008; Stanke et al., 2006) to generate putative gene models.

#### Functional annotation

Structural gene predictions were used as input for Eukaryotic Non-Model Transcriptome Annotation Pipeline (EnTAP; Hart et al., 2019), to add functional information and to and identify improbable gene models. EnTAP was run in runP mode with taxon = Acrogymnospermae using the RefSeq Plant and SwissProt databases plus a custom conifer protein database (O’Leary et al., 2016; The Uniprot Consortium, 2019). To further filter putative gene models, gFACs (Caballero and Wegrzyn, 2019) was used, first by separating multiexonic and monoexonic models.

Multiexonics were retained after filtering out models with non-canonical splice sites, micro-introns and micro-exons (<20 bp), and in-frame premature stop codons to ensure correct geneic structure. Additionally, to control for function, genes annotating through Inteproscan (Jones et. al., 2014) as retrodomains (including gag-polypeptide, retrotransposon, reverse transcriptase, copia, gypsy, and ty1) were discarded. In addition, any multi-exonic models that lacked functional annotation either with a sequence similarity hit or gene family assignment were removed. Additionally, gffcompare (https://ccb.jhu.edu/software/stringtie/gffcompare.shtml) identified overlap between gene models and softmasked regions of the genome, and multi-exonic gene models were removed if more than 50% of their length fell in masked regions. Clustered transcriptome sequences were aligned to the genome using GMAP (v. 2018-07-04; Wu & Watanabe, 2005; Wu & Nacu, 2010) with a minimum trimmed coverage of 0.95 and a minimum identity of 0.95. To determine overlap and nesting of gene models with this high confidence transcriptomic alignment, BEDtools (Quinlan and Hall, 2010). BUSCO v.3.0.2 (Simao et al., 2015) was used to assess the completeness of the filtered gene space.

#### Orthogroup assignment of proteins

Translated UniGenes for all available gymnosperms were downloaded from the forest genomics database TreeGenes (https://treegenesdb.org/; Wegrzyn et al., 2019; Falk et al., 2018). The corresponding files from the *Amborella trichopoda* genome assembly were also included to provide an outgroup to the gymnosperm taxa. Each taxon was evaluated for completeness with BUSCO v4.0.2 in protein mode. All taxa with over 60% completeness were included in OrthoFinder (Emms and Kelly 2015; Emms and Kelly 2019) to identify orthogroups. The longest sequence in each orthogroup was retained, regardless of source species. Species-specific orthogroups unique to giant sequoia were noted. The resulting nonredundant species-specific orthogroups were functionally annotated with EnTAP in runP mode with taxon = *Sequoiadendron* using the RefSeq Plant and SwissProt databases.

#### Gene family evolution

Following orthogroup assignment with OrthoFinder, a species tree and orthogroup statistics were used as input for CAFE v4.1 (Han et al., 2013) to assess gene family contraction and expansion dynamics, using a single birth/death parameter (λ) across the phylogeny. Gene families in the giant sequoia lineage experiencing rapid evolution were then functionally annotated using EnTAP.

#### Annotation and analysis of NLR genes

NLR genes were identified using the NLR-Annotator pipeline (Steuernagel et al., 2018) on the giant sequoia 2.0 assembly, then that output was cross-referenced with the genome annotation. Using the genome annotation file and the NLR gene file as input, the BEDtools intersect function (Quinlan and Hall, 2010) was used to identify putative NLRs that were also present in the annotation, requiring features in the NLR gene file to overlap with 100% of the annotation feature. NLR-gene maximum likelihood trees were generated with RAxML v8.2.12 (Stamatakis, 2014) using the amino acid sequence of the central NB-ARC [AB2] domain output by NLR-Annotator. NB-ARC domains that included greater than 50% missing data were excluded from all analyses. The best trees were visualized with the Interactive Tree of Life (iTOL) tool, with bootstrap values shown (Letunic and Bork, 2016). Determination of TIR and CC domains was based on motif data from Jupe and colleagues (2012). RPW8-like motifs were determined by alignment to a recently described RNL motif (CFLDLGxFP) (Van Ghelder et al., 2019).

## Data availability

The genome assembly of giant sequoia is available at NCBI under accession GCA_007115665.2, and raw sequence data are available under accessions SRX5827056 - SRX5827083. Annotation files are available at https://treegenesdb.org/FTP/Genomes/Segi.

## RESULTS AND DISCUSSION

### Sequencing and assembly

Assembly of the giant sequoia genome leveraged sequence data from four libraries (Table 1). Illumina reads (135x) from a haploid megagametophyte library combined with Oxford Nanopore sequence from foliage (21x) contributed to the contig assembly. The contig assembly was subsequently scaffolded with data from Dovetail Chicago (47x) and Hi-C libraries (76x) in succession.

**Table 1.** Data used for the giant sequoia assemblies from four library types.

|  |  |  |  |
| --- | --- | --- | --- |
| Type | Number of reads | Average read length (bp) | Estimated coverage |
| Illumina paired end | 7,752,481,576 | 2x151 | 135x |
| Oxford Nanopore MinION | 24,360,895 | 7,484 | 21x |
| Dovetail Chicago | 2,592,465,290 | 2x151 | 47x |
| Dovetail Hi-C | 4,202,954,328 | 2x151 | 76x |

#### Giant sequoia 1.0 assembly

Initial contig assembly of the Illumina and Oxford Nanopore sequence data yielded giant sequoia 1.0. The initial contig assembly giant sequoia 1.0 had a contig N50 of 359,531 bp and a scaffold N50 of 489,478.

Genome size was estimated by counting 31-mers (all sub-sequences of 31 bases) in the Illumina reads and computing the histogram of the kmer frequencies vs. counts using jellyfish tool version 2.0 (Marcais et al., 2011). The histogram of 31-mer frequency counts had its largest peak at 101 (see Figure 1). There was a small second peak at 204, roughly double the highest 31-mer frequency was 101, likely corresponding to 2x repeat sequences in the genome. The k-mer coverage of the genome was then estimated by computing the area under the curve for frequencies between 1 and 10000 and dividing that number by 101. This method arrived at the genome size estimate of 8,588 Gbp.

**Figure 1.**
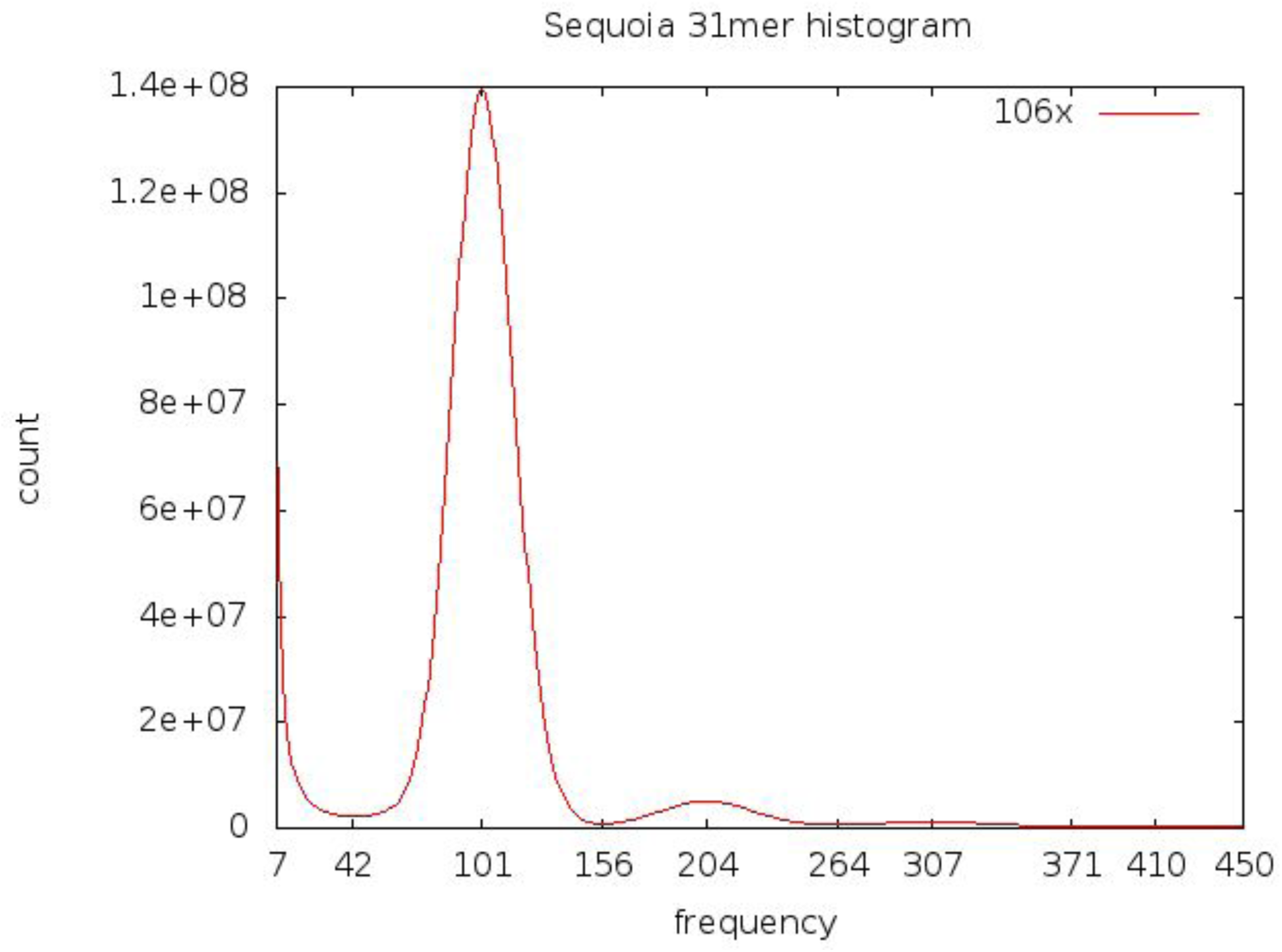
The histogram of 31-mer count in Illumina paired end reads. The red curve shows the number of 31-mers that are present in the reads X times, where X is the frequency plotted on horizontal axis. The main peak is at 101.

The intermediate step of correction of the Nanopore in MaSuRCA resulted in 24,279,305 mega-reads with an average read length of 6,726 bp. The consensus error rate for the assembly was estimated by aligning the Illumina reads to the contigs with bwa mem (Li, 2013) and then calling variants with freebayes (Garrison et al., 2012) software. Any site in the consensus that had no Illumina reads agreeing with the consensus and at least three Illumina reads agreeing on an alternative variant was considered an error. The total number of bases in the error variants were counted and divided by the total number of bases in the contigs. This yielded an assembly error rate of 0.3 errors per 10000 bases, or consensus quality of 99.997%.

The initial contig assembly giant sequoia 1.0 had a contig N50 of 347,954 bp and a scaffold N50 of 490,521.

#### Giant sequoia 2.0 assembly

The Dovetail HiRise Chicago and Hi-C assembly increased the total assembly size marginally, to 8.125 Gbp, but notably yielded a large increase in the N50 to 690.55Mb (Table 2). The overall number of scaffolds was reduced to 8,125, and the N90 of the final assembly was 690.55Mb. It is worth noting that the largest scaffold in this assembly is 985 Mbp in length, making it the longest contig assembled to date in any organism.

**Table 2.** Assembly statistics for the initial contig assembly giant sequoia 1.0 and the final scaffolded assembly giant sequoia 2.0.

| Assembly | Sequence (bp) | N50 contig | N50 scaffold | Number of contigs | Number of scaffolds |
| --- | --- | --- | --- | --- | --- |
| Giant sequoia 1.0 | 8,122,145,191 | 347,954 | 490,521 | 49,651 | 39,821 |
| Giant sequoia 2.0 | 8,125,622,286 | 347,954 | 690,549,816 | 52,886 | 8,125 |

The tandem repeat finder program (trf v4.09, G. Benson 1999) identified repeat elements up to 500 bp long, and those data were used to plot a histogram of repeat unit lengths which had peaks at 7, 181, and 359 bp. Based on the position and clustering along the chromosomes, the 7-mer was identified as the telomeric repeat and the 181-mer as the centromeric one.

The most common telomeric 7-mers were TTTAGGG (present in most land plants), and TTGAGGG. The two 7-mers alternate and have similar frequencies.

The 181 bp centromeric repeat unit consensus sequence was AAAAATTGGAGTTCGCGTGACACAGATGCAACGTAGCCTTAAAATCAGGTCTTCGCCGAACTCGACATTAAATCGATGGAAATTCAACATTCACGAAAACTGATAGAAAATAAAGGTTCTTAATAGTCATCTACAACACAATCTAAATCAAAGTTCTCCAAACATGGTTGATTATGGGTG.

By looking at the positions of the centromeric and telomeric repeats, a mis-assembly was identified in the original HiRise reference. Two centromeric and one telomeric region were located in the middle of the longest scaffold (1.82Gb), and subsequently this scaffold was split into chr1 (0.95Gb) and chr3 (0.84Gb).

There are 11 chromosomes in giant sequoia (Buccholz, 1939; later confirmed by Jensen and Levan, 1941 and Schlarbaum and Tschuiya, 1984), and the 11 largest scaffolds in the assembly span across the centromere (Table 3), suggesting a chromosome-level assembly. The 11 largest scaffolds range from 443 Mbp to 985 Mbp in size. Of these 11 scaffolds, seven include telomeric sequence on both ends. The remaining four scaffolds have telomeric sequence on one end. Beyond the 11 largest scaffolds, the next largest (Sc7zsyj_3574) (171 Mb) includes telomere at one end, suggesting it is a substantial portion of a chromosome arm for one of the scaffolds with only one telomere (chromosomes 1, 3, 6, and 9).

**Table 3.** Summary of largest scaffolds in giant sequoia 2.0 and presence of centromeric and telomeric repeat regions

| Scaffold ID | Length (bp) | Centromere? | Number of telomeres |
| --- | --- | --- | --- |
| chr1 | 986,618,365 | Y | 1 |
| chr2 | 873,713,311 | Y | 2 |
| chr3 | 843,110,718 | Y | 1 |
| chr4 | 722,823,090 | Y | 2 |
| chr5 | 690,549,816 | Y | 2 |
| chr6 | 676,903,824 | Y | 1 |
| chr7 | 659,235,867 | Y | 2 |
| chr8 | 649,867,199 | Y | 2 |
| chr9 | 641,211,466 | Y | 1 |
| chr10 | 632,191,860 | Y | 2 |
| chr11 | 443,565,592 | Y | 2 |
| Sc7zsyj_3574 | 171,454,409 | N | 1 |

#### Assessing assembly completeness

For a rough estimate of the assembly completeness, BUSCO v3.0.2 was run with the embryophyta database (Simao et al., 2015) of 1440 genes. For the complete giant sequoia 2.0 genome, the tool found 559 complete BUSCOs out of which 515 were in a single copy, 44 were duplicated, and 133 were fragmented BUSCOs (Table 4). Another 748 BUSCOs were missing. In both the full giant sequoia 2.0 assembly and the version filtered to remove all scaffolds smaller than 3 kbp, completeness was estimated at 38% using BUSCO. Assembly completeness of other conifer assemblies (Supplementary Table S1) range from 27-44%, suggesting giant sequoia 2.0 completeness is consistent with existing work. Despite the contiguity of the assembly, the BUSCO completeness of the genome appears lower than expected, likely due to the presence of very long introns in conifers, which can inhibit identification of genes.

**Table 4.** Completeness of assembly and gene sets assessed with BUSCOv3.0.2.

|  | **Giant sequoia v2.0** | **Giant sequoia v2.0**<br>**(≥3kbp)** | **Transcriptome** | **High-confidence**<br>**gene set** |
| --- | --- | --- | --- | --- |
| Number of input sequences | 8215 | 8120 | 25859 | 37936 |
| Complete BUSCOs (C) | 559 | 553 | 1139 | 766 |
| Complete and single-copy<br>BUSCOs (S) | 515 | 508 | 1076 | 683 |
| Complete and duplicated<br>BUSCOs (D) | 44 | 45 | 63 | 83 |
| Fragmented BUSCOs (F) | 133 | 131 | 66 | 149 |
| Missing BUSCOs (M) | 748 | 756 | 235 | 525 |
| Total BUSCO groups<br>searched | 1440 | 1440 | 1440 | 1440 |
| Percentage found | 38.82% | 38.40% | 79.10% | 53.19% |

#### Comparison to existing gymnosperm assemblies

The contiguity of giant sequoia 2.0 is most apparent when comparing with other gymnosperm assemblies (Table 5). Giant sequoia 2.0 has an N50 scaffold size of 690Mb, an order of magnitude larger than scaffold N50s reported in other conifers.

**Table 5.** Comparison of giant sequoia v2.0 assembly and annotation to selected gymnosperm genome projects. 5a shows assembly statistics as reported in referenced manuscripts. 5b shows annotation statistics as calculated using gFACs on most recent annotations available at TreeGenes. Annotation statistics for *Picea glauca* are reported as in referenced manuscript.

| 5a | *Sequoiadendron giganteum* | *Abies alba* | *Picea glauca* | *Pinus lambertiana* | *Pinus taeda* | *Pseudotsuga menzesii* | *Ginkgo biloba* | *Gnetum montanum* |
| --- | --- | --- | --- | --- | --- | --- | --- | --- |
| Reference |  | Mosca et al., 2019 | Warren et al., 2015 | Stevens et al., 2016 | Neale et al., 2014 | Neale et al., 2017 | Guan et al., 2016 | Wan et al., 2018 |
| Genome size (Mbp) | 8,114 | 18,167 | 20,000 | 31,000 | 20,613 | 15,700 | 10,610 | 4,110 |
| Chromosomes | 11 | 12 | 12 | 12 | 12 | 12 | 12 | 22 |
| TE content (%) | 79 | 78 | N/A | 79 | 81 | 72 | 77 | 86 |
| N50 scaffold size (kb) | 690,549.82 | 14.05 | 71.50 | 246.60 | 107.04 | 340.70 | 1,360.00 | 475.17 |
| 5b | *Sequoiadendron giganteum* | *Abies alba* | *Picea glauca* | *Pinus lambertiana* | *Pinus taeda* | *Pseudotsuga menzesii* | *Ginkgo biloba* | *Gnetum montanum* |
| Number of genes | 37,936 | 94,209 | 14,462 | 38,518 | 51,751 | 46,688 | 41,840 | 27,493 |
| Average overall CDS size | 1,084 | 629 | 1,421 | 1,102 | 1,131 | 1,180 | 1,186 | 1,290 |
| Average size multiexonic introns | 4,067 | 315 | 603 | 11,468 | 5,596 | 4,685 | 7,884 | 1,769 |
| Maximum intron length (kb) | 1,399.11 | 36.01 | 119.32 | 1,254.69 | 758.52 | 351.90 | 1,272.92 | 342.13 |

### Annotation of giant sequoia 2.0

#### Repeat annotation

Using the custom repeat database created by RepeatModeler, the majority (72.85%) of the giant sequoia genome was softmasked. Subsequent masking using conifer-specific repeat libraries yielded an additional 6% of masked sequence. LTRs were the most abundant known element (28%, Supplementary Table S2) in the masked sequence. These results are comparable to observations from different conifer species, e.g. the most recent *Pinus lambertiana* assembly contained 79% repetitive sequence (Stevens et al., 2016). That our observations are consistent with the only conifer lineage sequenced until now (Pinaceae) is not surprising, as all conifers have large genome sizes, and this genomic bloat is attributed to the proliferation of repetitive elements throughout the genome (Neale et al., 2014).

#### Gene Annotation

Structural annotation using BRAKER2 resulted in 1,460,545 predicted gene models, with an average intron length of 2,362 bp (Table 6). The average CDS length was 613 bp, including both multi- and mono-exonic models. The initial gene set included models with long introns, with the longest measuring 385,133 bp. The number of mono-exonic genes (941,659) was almost twice as large as the total number of multi-exonic gene models (518,886). Even with reasonable filters, the number of *ab initio* prediction of mono-exonic genes was highly inflated. Therefore, the mono-exonic *ab initio* genes were removed from the gene space. The *ab initio* gene space was expanded by the addition of 14,538 well aligned unique transcriptome sequences of which 6,982 are mono-exonic and the remaining 7,556 are multi-exonic. After filtering, annotation yielded 37,936 high quality gene models. The average CDS length increased to 1,083 bp. The proportion of mono-exonics (5,163) to multi-exonics (32,773) was drastically reduced using the transcriptome as an evidence source. Long introns were maintained, with the max intron length in the high quality set reaching nearly 1.4 Mb.

**Table 6.** Gene models proposed by BRAKER2, before and after filtering. Intermediate set was filtered by removing monoexonic models, models with greater than 50% of their length in a masked region, models annotated as retrodomains, and models lacking functional annotation with EnTAP. The high-confidence set includes the intermediate set, plus mono- and multi-exonic models derived from transcript evidence, removing any fully nested gene models.

|  | **Initial**<br>**model set** | **Intermediate**<br>**filtered set** | **High-confidence**<br>**set** |
| --- | --- | --- | --- |
| Total Genes | 1,460,545 | 32,360 | 37,936 |
| Average CDS length | 613.90 | 1099.08 | 1083.00 |
| Average number of exons | 2.78 | 4.22 | 4.5 |
| Average intron length (bp) | 2,362 | 2,233 | 4,066 |
| Max intron length | 385,133 | 159,979 | 1,399,110 |
| Total monoexonics | 941,659 | - | 5,163 |
| Total multiexonics | 518,886 | 32,360 | 32,773 |

Of the 37,936 high quality gene models, 35,183 were functionally annotated by either sequence similarity search or gene family assignment with EnTAP. These functionally annotated gene models include the longest plant intron found so far, at 1.4 Mb. Large introns are characteristic of conifer genomes, with introns up to 800 Kbp observed in *Pinus taeda* (Wegrzyn et al., 2014) and introns over 500 Kbp in *Pinus lambertiana* (Stevens et al., 2016).

Functional annotation of the gene containing the 1.4 Mb long intron suggests it is a member of the WASP (Wiskott-Aldrich syndrome protein) family. Wiskott-Aldrich syndrome proteins are in turn members of the SCAR/WAVE (suppressor of cAMP receptor/WASP family verprolin homologous) gene regulatory complex, which in plants has an important role in cell morphogenesis via activation of actin filament proteins (Yanagisawa, Zhang, and Szymanski, 2013).

Distribution of the high-quality gene models spanned the length of all 11 chromosomes (Figure 2). Repeat density varied across the chromosomes, including overlap with annotated regions.

**Figure 2.**
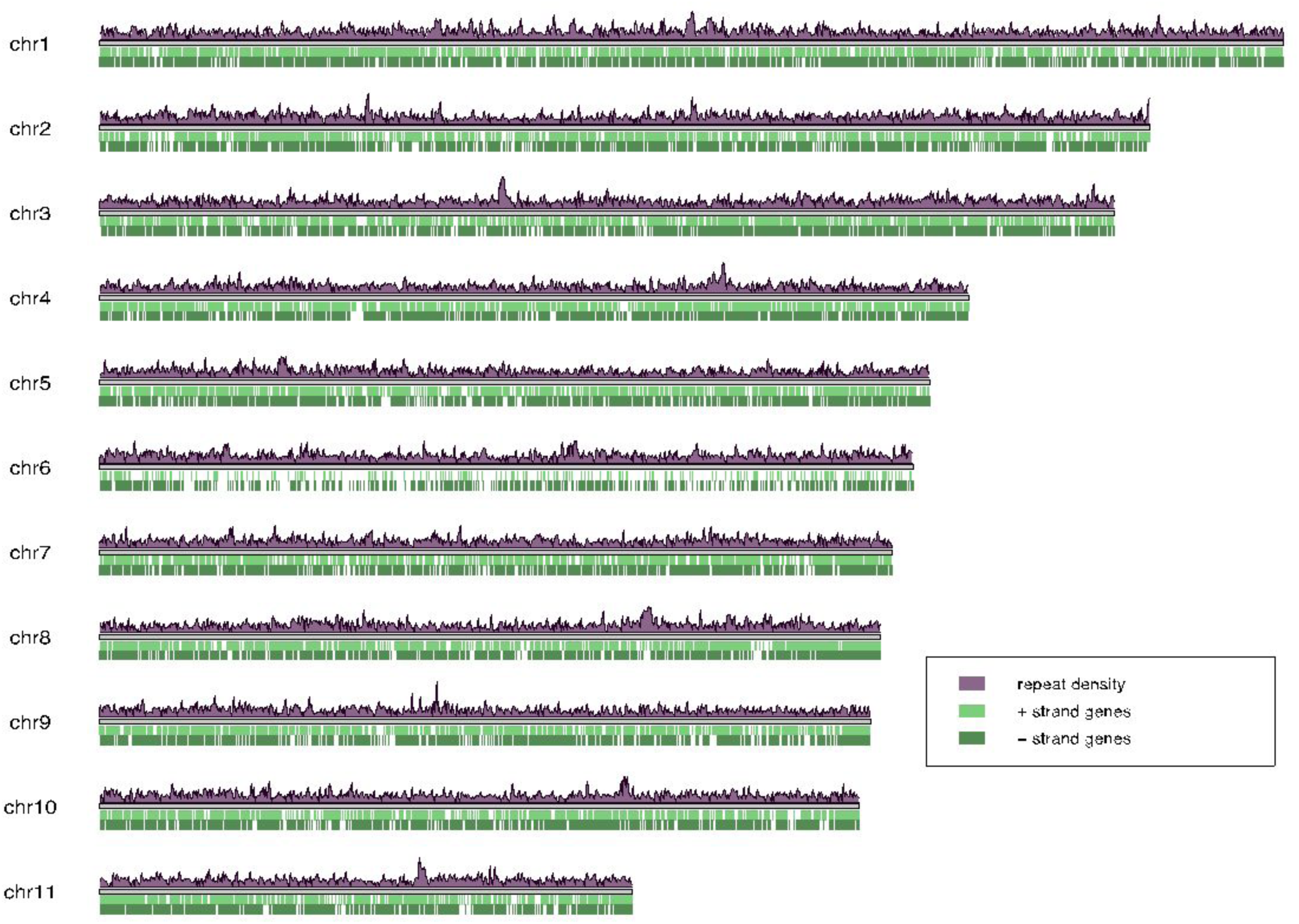
Repeat density and gene content of giant sequoia 2.0. Light green bars are + strand genes, dark green bars are - strand genes. Repeat density in purple, plotted in 1kb windows.

#### Assessing annotation completeness

Completeness of the annotation was assessed with BUSCO (Table 4). The independent transcriptome completeness of 79% represents the maximum possible BUSCO score for the gene model sets. The BUSCO completeness of the final high-quality gene set was 53%, comparable to the same metric in *Pinus taeda* (53%, Wegrzyn et al., 2014) and *Pinus lambertiana* (50%, Stevens et al., 2016), suggesting the annotation of giant sequoia is on par with other conifer genomes.

##### Comparison to existing gymnosperm annotations

While the genome size of giant sequoia is rather small for a gymnosperm (Table 5), the identified repeat content of giant sequoia 2.0 (79%) is in line with observations from other taxa. The number of high quality annotated genes (37,936) is higher than many gymnosperm assemblies, though there is substantial variation in annotation results across the lineage. Average CDS length and average intron length in giant sequoia 2.0 fall within the observed ranges for existing assemblies, though notably the longest intron reported here is ∼1.4 Mb, nearly 400kb longer than the previous longest intron (from *Pinus taeda*, at over 800 kbp). That giant sequoia 2.0 contains an even longer intron is likely due to the contiguity of our assembly, which is unprecedented in conifers.

#### Orthology assignment and gene family evolution

Using unigene sets from TreeGenes, twenty gymnosperm taxa passed the 60% threshold for BUSCO completeness (Table 7). Orthogroup clustering of 695,700 protein sequences from these twenty gymnosperms plus an outgroup (*Amborella trichopoda*) yielded a total of 44,797 orthogroups (Supplementary Table S3). Only 206 were single-copy in all species, and 5,953 orthogroups had representatives from each species. Overall, 6.5% of all protein sequences were in species-specific orthogroups. Of the species-specific orthogroups (12,121 in total), 607 were unique to giant sequoia (Table 8). Among the 607 giant sequoia-specific orthogroups, 536 were functionally annotated with either gene family assignment (318) sequence similarity search (8) or both (536) (Supplementary Table S4).

**Table 7.** BUSCO completeness for 20 gymnosperm taxa and an angiosperm outgroup (*Amborella trichopoda*)

| TreeGenes code | Abba | Gibi | Gnmo | Megl | Pama | Pial | Piba |
| --- | --- | --- | --- | --- | --- | --- | --- |
| taxon | *Abies balsamea* | *Ginkgo biloba* | *Gnetum gnemon* | *Metasequoia glyptostroboides* | *Picea mariana* | *Pinus albicaulis* | *Pinus banksiana* |
|  | Balsam fir | Ginkgo | Gnemon/milinjo | Dawn redwood | Black spruce | White pine | Jack pine |
| Unigene set |  |  |  |  |  |  |  |
| Data source(s) | transcriptome | annotation,<br>transcriptome | annotation | transcriptome | transcriptome | transcriptome | transcriptome |
| Number of unigenes | 21,250 | 110,296 | 21,887 | 19,237 | 22,876 | 27,226 | 21,278 |
| Average length of unigenes | 396.46 | 269.39 | 351.69 | 343.17 | 376.93 | 338.76 | 381.58 |
| **BUSCOv4.0.2** |  |  |  |  |  |  |  |
| Complete | 1419 | 1437 | 1301 | 1109 | 1429 | 1453 | 1342 |
| Complete & single copy | 1357 | 1292 | 1265 | 1068 | 1377 | 1398 | 1283 |
| Complete & duplicated | 62 | 145 | 36 | 41 | 52 | 55 | 59 |
| Fragmented | 52 | 89 | 82 | 206 | 66 | 48 | 93 |
| Missing | 143 | 88 | 231 | 299 | 119 | 113 | 179 |
| Total searched | 1614 | 1614 | 1614 | 1614 | 1614 | 1614 | 1614 |
| % complete | 87.92% | 89.03% | 80.61% | 68.71% | 88.54% | 90.02% | 83.15% |

| TreeGenes code | Pice | Picn | Pila | Pima | Pimn | Pipt | Pist |
| --- | --- | --- | --- | --- | --- | --- | --- |
| taxon | *Pinus cembra* | *Pinus canariensis* | *Pinus lambertiana* | *Pinus massoniana* | *Pinus monticola* | *Pinus patula* | *Pinus strobus* |
|  | Swiss stone pine | Canary island pine | Sugar pine | Chinese red pine | Western white pine | Mexican weeping pine | Eastern white pine |
| Unigene set |  |  |  |  |  |  |  |
| Data source(s) | transcriptome | transcriptome | annotation,<br>transcriptome | transcriptome | transcriptome | transcriptome | transcriptome |
| Number of unigenes | 17,994 | 22,631 | 42,256 | 33,891 | 17,447 | 46,563 | 21,697 |
| Average length of unigenes | 411.38 | 327.27 | 357.80 | 322.13 | 388.92 | 348.04 | 372.89 |
| **BUSCOv4.0.2** |  |  |  |  |  |  |  |
| Complete | 1300 | 1183 | 1369 | 1415 | 1202 | 1526 | 1338 |
| Complete & single copy | 1250 | 1147 | 1276 | 1367 | 1163 | 1435 | 1296 |
| Complete & duplicated | 50 | 36 | 93 | 48 | 39 | 91 | 42 |
| Fragmented | 119 | 226 | 88 | 84 | 146 | 28 | 95 |
| Missing | 195 | 205 | 157 | 115 | 266 | 60 | 181 |
| Total searched | 1614 | 1614 | 1614 | 1614 | 1614 | 1614 | 1614 |
| % complete | 80.55% | 73.30% | 84.82% | 87.67% | 74.47% | 94.55% | 82.90% |

| TreeGenes code | Pita | Pnte | Psme | Segi | Sese | Thoc | Amtr |
| --- | --- | --- | --- | --- | --- | --- | --- |
| taxon | *Pinus taeda* | *Pinus tecunumanii* | *Pseudotsuga menziesii* | *Sequoiadendron giganteum* | *Sequoia sempervirens* | *Thuja occidentalis* | *Amborella trichopoda* |
|  | Loblolly pine | Tecun Uman Pine | Douglas-fir | Giant sequoia | Coast redwood | Eastern white cedar |  |
| Unigene set |  |  |  |  |  |  |  |
| Data source(s) | annotation, transcriptome | transcriptome | annotation, transcriptome | annotation | transcriptome | transcriptome | annotation |
| Number of unigenes | 45255 | 22287 | 70036 | 42325 | 21798 | 19208 | 24753 |
| Average length of unigenes | 392.03 | 450.75 | 289.17 | 328.80 | 303.28 | 338.63 | 318.60 |
| BUSCOv4.0.2 |  |  |  |  |  |  |  |
| Complete | 1090 | 1517 | 1152 | 1113 | 1064 | 1187 | 1303 |
| Complete & single copy | 1000 | 1453 | 1030 | 1057 | 1030 | 1149 | 1292 |
| Complete & duplicated | 90 | 64 | 122 | 56 | 34 | 38 | 11 |
| Fragmented | 161 | 22 | 253 | 232 | 221 | 172 | 49 |
| Missing | 363 | 75 | 209 | 269 | 329 | 255 | 23 |
| Total searched | 1614 | 1614 | 1614 | 1614 | 1614 | 1614 | 1614 |
| % complete | 67.53% | 93.99% | 71.38% | 68.96% | 65.92% | 73.54% | 80.73% |

\*Not a TreeGenes code; Amtr peptide data were downloaded from Ensembl (Howe et al., 2019).

**Table 8.**
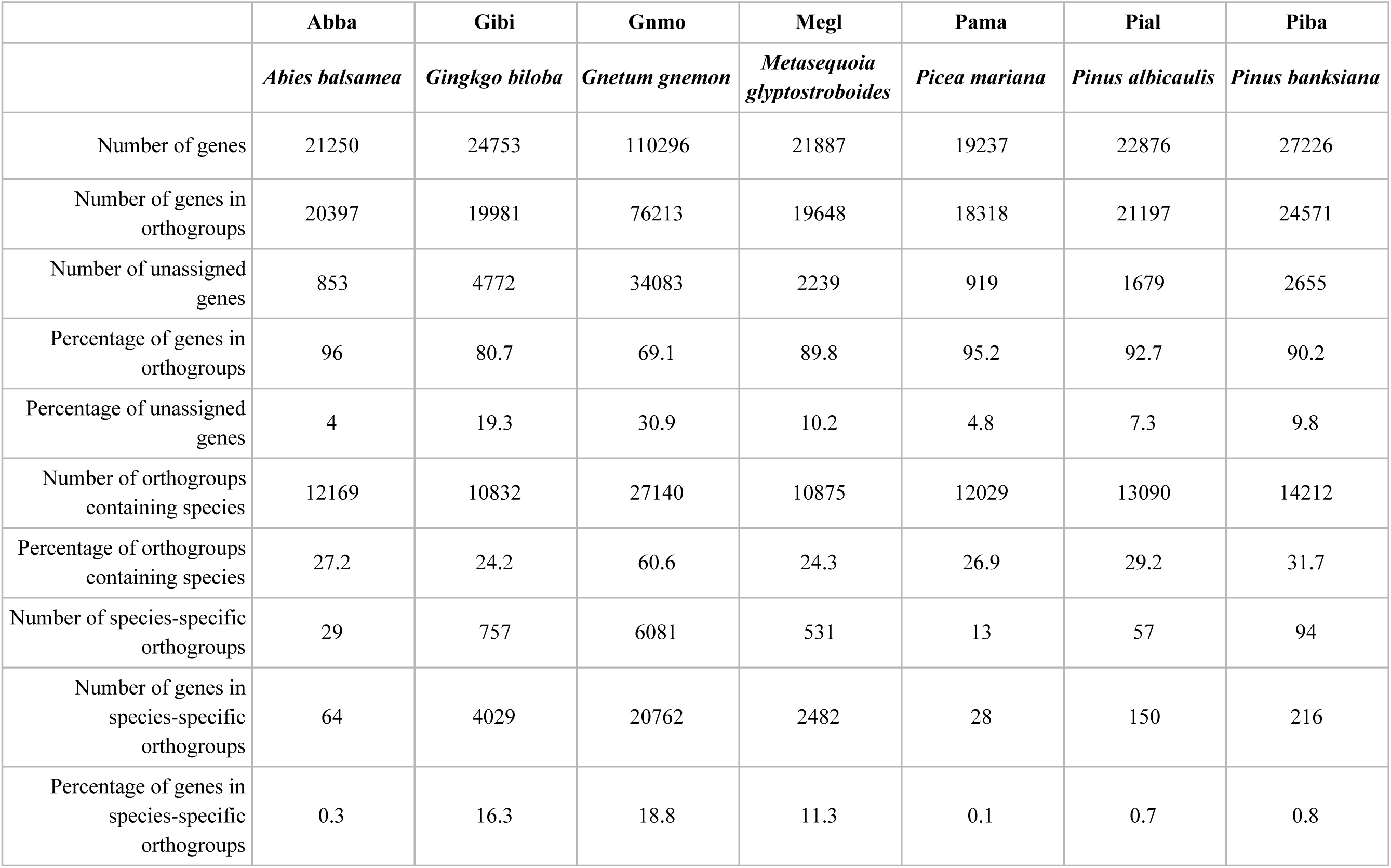
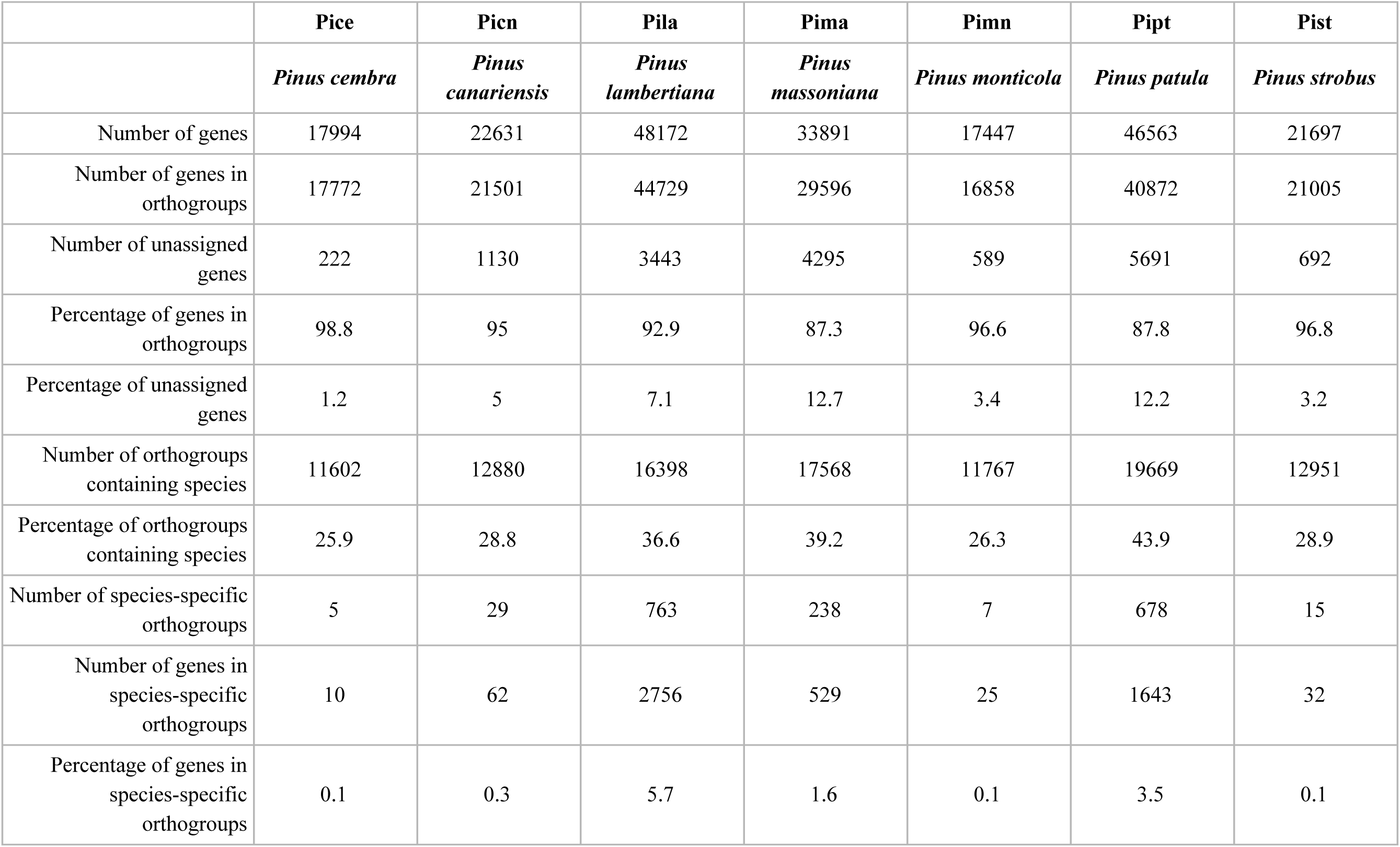
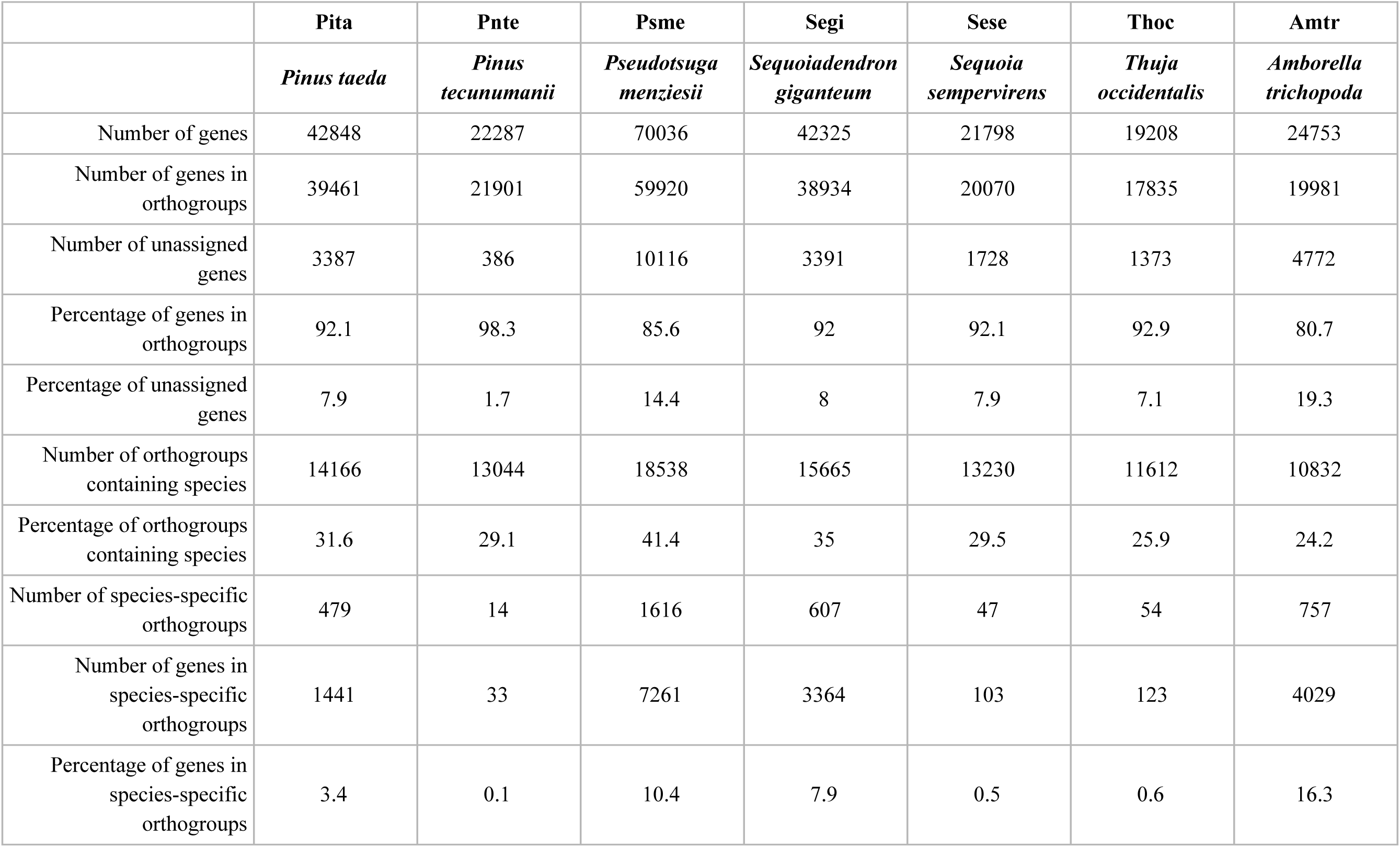
Orthogroup assignment summary for 20 gymnosperm taxa and an angiosperm outgroup (*Amborella trichopoda*; Amtr).

|  | **Abba** | **Gibi** | **Gnmo** | **Megl** | **Pama** | **Pial** | **Piba** |
| --- | --- | --- | --- | --- | --- | --- | --- |
|  | *Abies balsamea* | *Gingko biloba* | *Gnetum gnemon* | *Metasequoia glyptostroboides* | *Picea mariana* | *Pinus albicaulis* | *Pinus banksiana* |
| Number of genes | 21250 | 24753 | 110296 | 21887 | 19237 | 22876 | 27226 |
| Number of genes in orthogroups | 20397 | 19981 | 76213 | 19648 | 18318 | 21197 | 24571 |
| Number of unassigned genes | 853 | 4772 | 34083 | 2239 | 919 | 1679 | 2655 |
| Percentage of genes in orthogroups | 96 | 80.7 | 69.1 | 89.8 | 95.2 | 92.7 | 90.2 |
| Percentage of unassigned genes | 4 | 19.3 | 30.9 | 10.2 | 4.8 | 7.3 | 9.8 |
| Number of orthogroups containing species | 12169 | 10832 | 27140 | 10875 | 12029 | 13090 | 14212 |
| Percentage of orthogroups containing species | 27.2 | 24.2 | 60.6 | 24.3 | 26.9 | 29.2 | 31.7 |
| Number of species-specific orthogroups | 29 | 757 | 6081 | 531 | 13 | 57 | 94 |
| Number of genes in species-specific orthogroups | 64 | 4029 | 20762 | 2482 | 28 | 150 | 216 |
| Percentage of genes in species-specific orthogroups | 0.3 | 16.3 | 18.8 | 11.3 | 0.1 | 0.7 | 0.8 |

|  | **Pice** | **Picn** | **Pila** | **Pima** | **Pimn** | **Pipt** | **Pist** |
| --- | --- | --- | --- | --- | --- | --- | --- |
|  | *Pinus cembra* | *Pinus canariensis* | *Pinus lambertiana* | *Pinus massoniana* | *Pinus monticola* | *Pinus patula* | *Pinus strobus* |
| Number of genes | 17994 | 22631 | 48172 | 33891 | 17447 | 46563 | 21697 |
| Number of genes in orthogroups | 17772 | 21501 | 44729 | 29596 | 16858 | 40872 | 21005 |
| Number of unassigned genes | 222 | 1130 | 3443 | 4295 | 589 | 5691 | 692 |
| Percentage of genes in orthogroups | 98.8 | 95 | 92.9 | 87.3 | 96.6 | 87.8 | 96.8 |
| Percentage of unassigned genes | 1.2 | 5 | 7.1 | 12.7 | 3.4 | 12.2 | 3.2 |
| Number of orthogroups containing species | 11602 | 12880 | 16398 | 17568 | 11767 | 19669 | 12951 |
| Percentage of orthogroups containing species | 25.9 | 28.8 | 36.6 | 39.2 | 26.3 | 43.9 | 28.9 |
| Number of species-specific orthogroups | 5 | 29 | 763 | 238 | 7 | 678 | 15 |
| Number of genes in species-specific orthogroups | 10 | 62 | 2756 | 529 | 25 | 1643 | 32 |
| Percentage of genes in species-specific orthogroups | 0.1 | 0.3 | 5.7 | 1.6 | 0.1 | 3.5 | 0.1 |

|  | **Pita** | **Pnte** | **Psme** | **Segi** | **Sese** | **Thoc** | **Amtr** |
| --- | --- | --- | --- | --- | --- | --- | --- |
|  | *Pinus taeda* | *Pinus tecunumanii* | *Pseudotsuga menziesii* | *Sequoiadendron giganteum* | *Sequoia sempervirens* | *Thuja occidentalis* | *Amborella trichopoda* |
| Number of genes | 42848 | 22287 | 70036 | 42325 | 21798 | 19208 | 24753 |
| Number of genes in orthogroups | 39461 | 21901 | 59920 | 38934 | 20070 | 17835 | 19981 |
| Number of unassigned genes | 3387 | 386 | 10116 | 3391 | 1728 | 1373 | 4772 |
| Percentage of genes in orthogroups | 92.1 | 98.3 | 85.6 | 92 | 92.1 | 92.9 | 80.7 |
| Percentage of unassigned genes | 7.9 | 1.7 | 14.4 | 8 | 7.9 | 7.1 | 19.3 |
| Number of orthogroups containing species | 14166 | 13044 | 18538 | 15665 | 13230 | 11612 | 10832 |
| Percentage of orthogroups containing species | 31.6 | 29.1 | 41.4 | 35 | 29.5 | 25.9 | 24.2 |
| Number of species-specific orthogroups | 479 | 14 | 1616 | 607 | 47 | 54 | 757 |
| Number of genes in species-specific orthogroups | 1441 | 33 | 7261 | 3364 | 103 | 123 | 4029 |
| Percentage of genes in species-specific orthogroups | 3.4 | 0.1 | 10.4 | 7.9 | 0.5 | 0.6 | 16.3 |

Orthogroup assignments were used as branch labels on a rooted species tree to show gene family contraction and expansion. On branch is the number of families that experienced expansion (dark blue, above) or contraction (light blue, below) (see Figure 3). Giant sequoia (Segi) experienced an overall expansion, with 4,953 families expanding and 1,923 families contracting since the species last shared common ancestor with coast redwood (*Sequoia sempervirens*; Sese).

**Figure 3:**
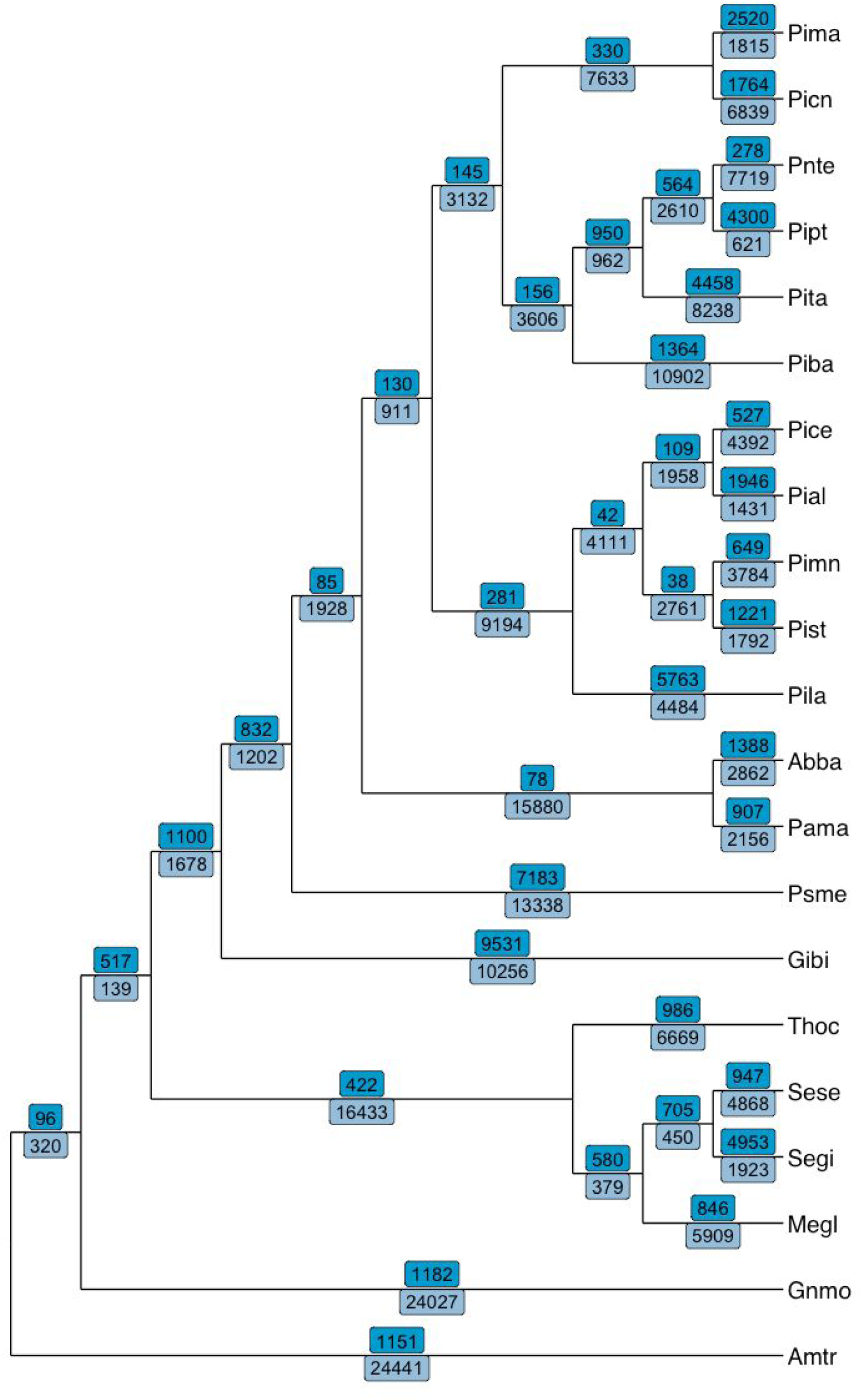
Gene family evolution along a gymnosperm cladogram. Numbers of expanded (bright blue, above branches) and contracted (light blue, below branches) orthogroups indicated in along each branch. Giant sequoia (Segi) experienced an overall expansion, with 4,953 orthogroups expanding and 1,923 contracting.

The expansions and contractions were further examined to identify nodes that experienced particularly rapid evolution. Many representatives of the Pinaceae have thousands of gene families that experienced rapid evolution since their lineages diverged (Figure 4). Along the branch to giant sequoia (Segi), 4,176 orthologous groups evolved rapidly. The majority of these 4,176 orthogroups are moderately represented in the giant sequoia dataset (e.g. with two to four members in an orthogroup), while others contain dozens of paralogs, up to over a hundred orthogroup members. Extracting the longest sequence from each of these yielded functional annotation with EnTAP for 3,994 of the rapidly evolving orthogroups. Rapidly expanding families were associated with primarily metabolic processes (GO:0090304, GO:0006796, GO:0044267) and macromolecule synthesis(GO:0009059, GO:0034645), in addition to molecular functions including metal-ion binding (GO:0046872), purine nucleotide (GO:0017076) and nucleoside (GO:0001883) binding, and kinase activity (GO:0016301). Rapidly contracting families were associated with biological processes such as protein (GO:0036211) and macromolecule modification (GO:0043412 and metabolic processes (GO:0044267, GO:0006796), and molecular functions including purine binding with nucleotides (GO:0017076) and nucleosides (GO:0001883), and phosphotransferase activity (GO:0016773).

**Figure 4:**
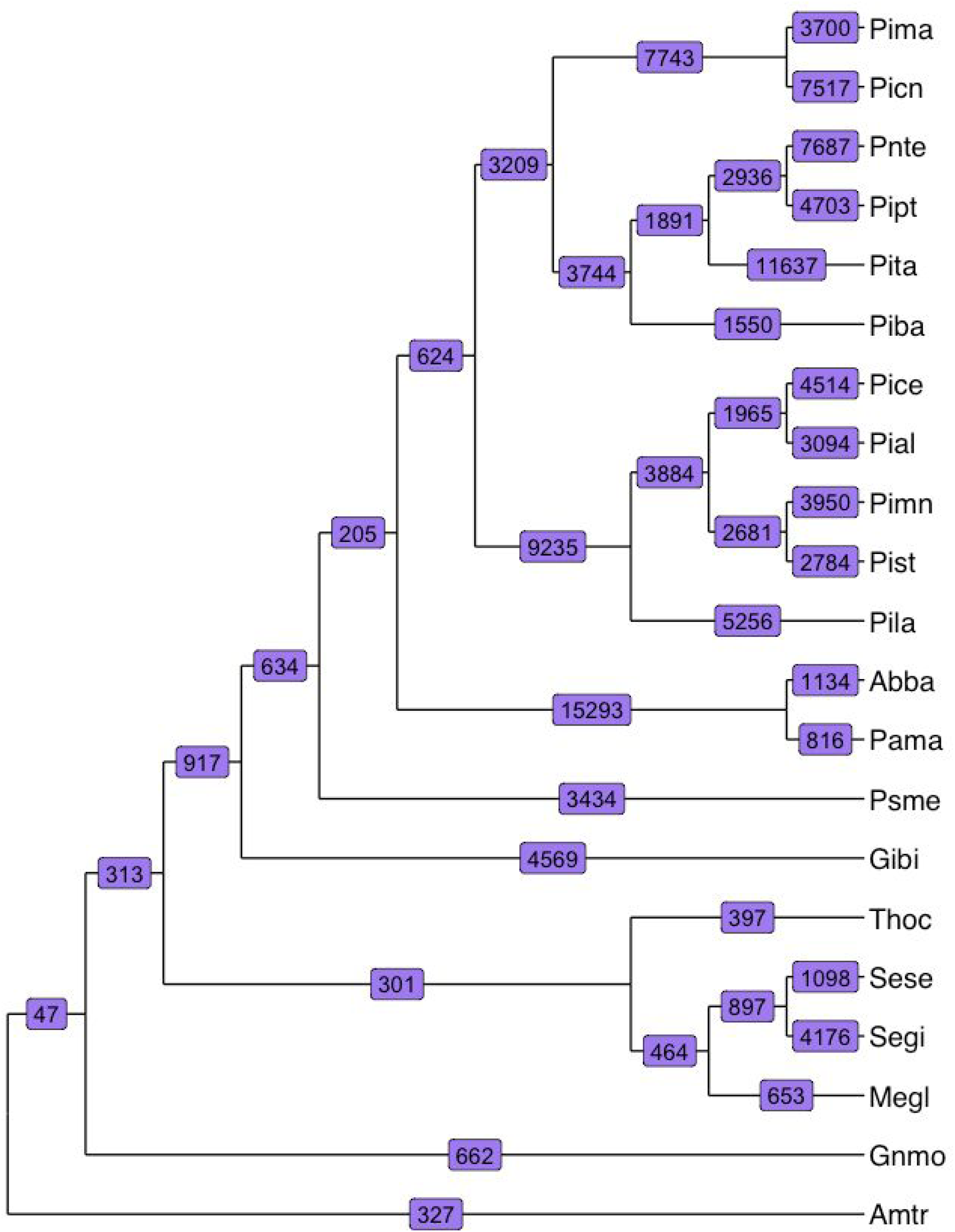
Rapid evolution along a gymnosperm cladogram. Numbers on each branch indicate the number of rapidly evolving gene families. Giant sequoia (Segi) has experienced rapid evolution in 4,176 gene families.

#### NLR genes in the giant sequoia genome

NLR proteins are structurally modular, typically containing an N-terminal coiled-coil (CC) domain, a Toll/interleukin-1 receptor (TIR) domain, or more rarely an RPW8-like CC domain; a conserved nucleotide binding domain (NB-ARC); and a C-terminal region comprising a variable number of leucine-rich repeats (LRRs) (Monteiro and Nishimura, 2018). NLR genes in giant sequoia 2.0 were identified by first running the genomic sequence through the NLR-Annotator pipeline (Steuernagel et al., 2018). Importantly, this pipeline does not require masking of repetitive regions and does not rely on gene model predictions. NLR-Annotator outputs are categorized as either ‘complete’ or ‘partial’ depending on whether all canonical domains (CC/TIR, NB-ARC, LRR) are present, and then further categorized as ‘pseudo-’ if a stop codon is predicted in any domain. All categorizations should be interpreted with care because the NLR-Annotator algorithm does not take intron/exon boundaries into account.

A total of 984 NLR genes were predicted by NLR-Annotator, of which 442 were identified as complete, 332 complete pseudo-, 88 partial, and 122 partial pseudo-. Seven hundred and twelve included intact NB-ARC domains with fewer than 50% gaps in the alignment. NLR-gene coordinates of all NLR gene sequences, and the relationships of the 712 based on an NB-ARC domain maximum likelihood tree are included in Supplementary Table S5, S6, and S7 as well as Supplementary Figure S1 . This number is roughly twice the number found in cultivated rice (Zhou et al., 2004; Read et al., 2020) and is consistent with other conifers (Van Ghelder et al., 2019).

NLR-Annotator identifies all suspected NLR motif-encoding regions of the genome. This likely includes true pseudogenes or gene fragments, both of which are important from an evolutionary perspective, but do not reflect the functional NLR arsenal. The NLR-Annotator output was cross-referenced with the giant sequoia genome annotation to identify the NLR genes that are supported by the annotation and therefore likely part of this arsenal; we refer to these 315 genes as consensus NLR genes. Of these, 211 were categorized by NLR-Annotator as complete, 65 as complete pseudo-, 29 as partial, and 10 as partial pseudo-. Two hundred and fifty seven of the 315 consensus NLR genes encode NB-ARC domains that met our criteria (see Methods); a maximum likelihood tree was generated using these domains (Figure 5). Coordinates of the genes and their NB-ARC sequences are included in Supplementary Table S5 and S7. NLR-Annotator predicted, non-consensus NLR genes may represent genes missed by the annotation, pseudogenes, or false positives.

**Figure 5:**
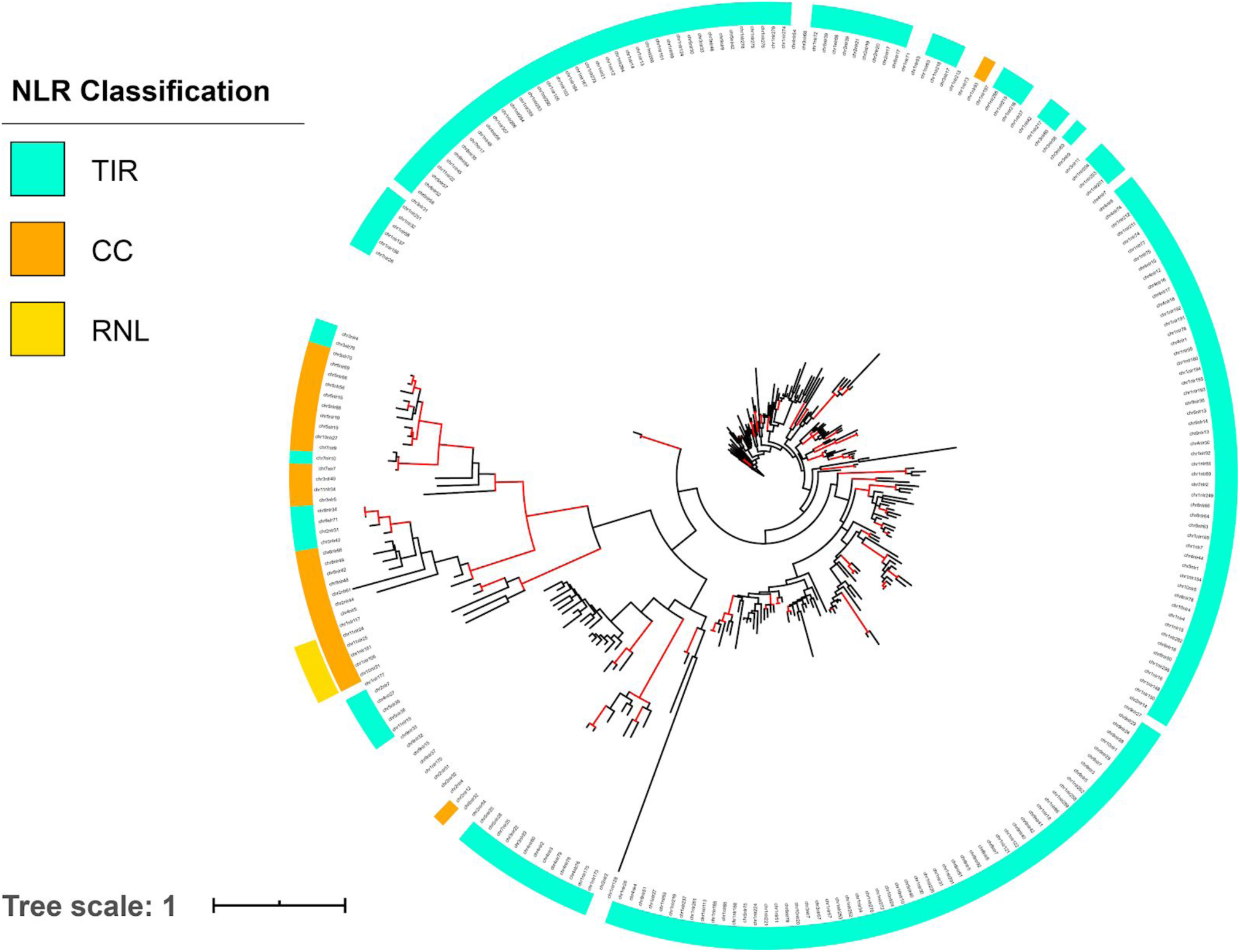
Maximum likelihood tree of NB-ARC domains of the 257 consensus NLR genes detected in the Segi assembly. Red branches indicate bootstrap support greater than 70%. The inner ring indicates predicted N-terminal TIR (blue) or CC (orange) domains. The outer ring indicates presence of an RPW8 motif present in the RNL sub-group of CC-NLRs. Tree is available at: http://itol.embl.de/shared/acr242

To investigate the evolution of NLR genes in giant sequoia, the list of consensus NLRs was compared with orthogroup assignments. Overall, consensus NLRs had membership in 63 orthogroups. Assessing the change in orthogroup size along each branch of the phylogeny revealed rapid expansion in NLR-associated orthogroups across the tree (Figure 6). Along the branch leading to giant sequoia (Segi), 34 NLR orthogroups expanded rapidly. The shared ancestors of giant sequoia and its closest relative, coast redwood (Sese), experienced rapid expansion in 11 NLR orthogroups. After the divergence of the California redwoods, five additional NLR orthogroups rapidly expanded in coast redwood, compared to the 34 rapidly expanding NLR orthogroups in giant sequoia. This pattern, a larger number of NLR orthogroups rapidly expanding in giant sequoia compared to coast redwood, is consistent with the numbers of all rapidly evolving orthogroups in each lineage (Figure 4).

**Figure 6:**
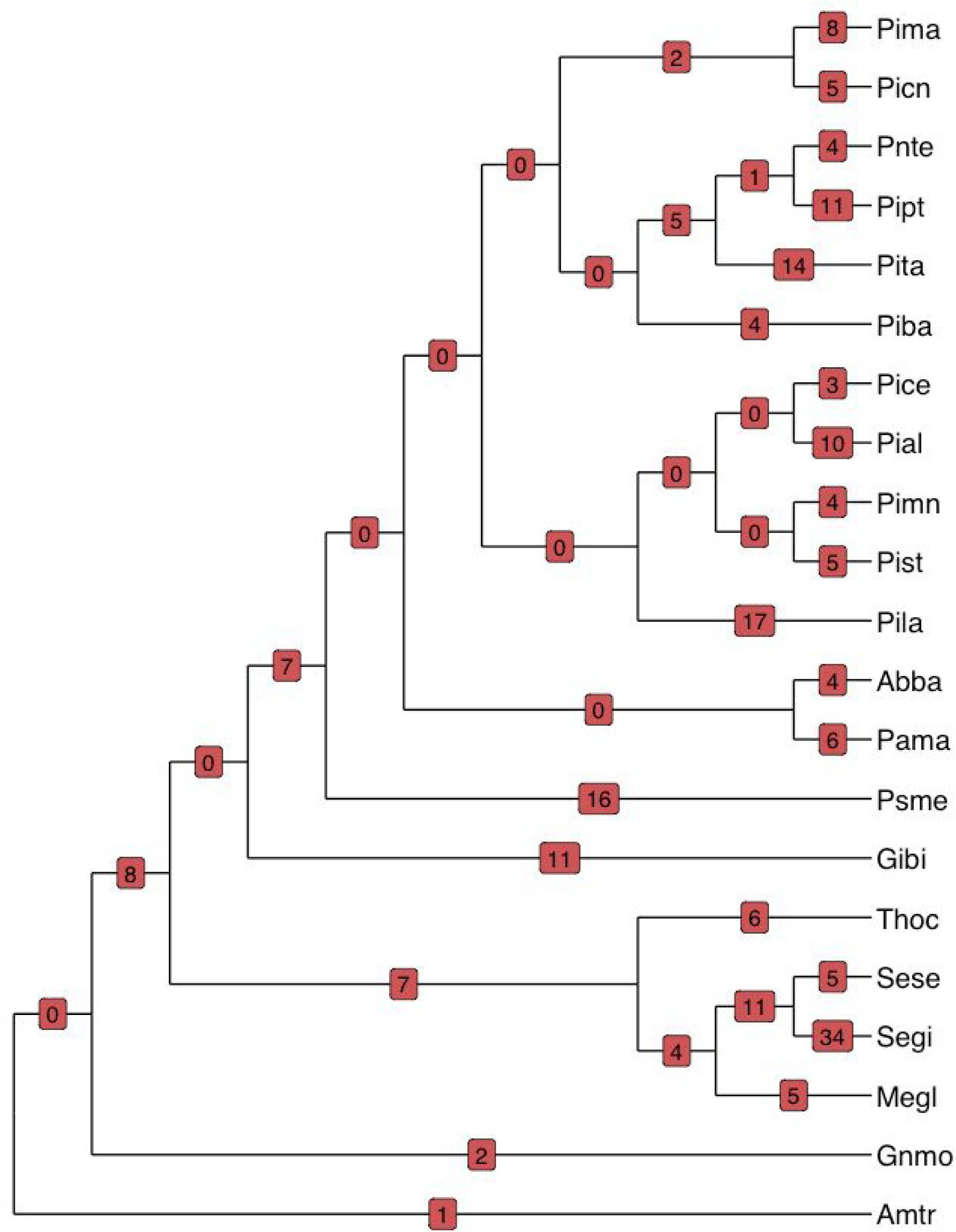
Rapid expansion in NLR-associated orthogroups along a gymnosperm cladogram. Numbers (red) on each branch indicate the number of rapidly expanding NLR orthogroups. Giant sequoia (Segi) has experienced rapid expansion in 34 NLR-associated orthogroups.

While the shared and unique NLR orthogroups identified in giant sequoia, coast redwood, and their common ancestors are perhaps associated with the observed pest resilience in both species, further work will be necessary to fully characterize the evolutionary patterns and functional roles of NLR gene families in redwoods and conifers as a whole.

## SUMMARY AND CONCLUSIONS

The high quality of this assembly demonstrates the value of combining multiple sequencing technologies and leveraging a unique biological feature of conifers (sufficient haploid megagametophyte tissue for sequencing), along with the value of incorporating chromosome-conformation capture libraries to allow improvements in scaffolding. The giant sequoia genome assembly presented here provides a robust foundation for ongoing genomic studies to identify groves with evidence of local adaptation, with a focus on not only NLR genes but the many other genes and gene families potentially useful in conservation and management.

For the future, inferences about the evolutionary trajectory of conifers (and gymnosperms) will require a broadening of taxonomic focus. As the vast majority of conifer genomic research is centered on Pinaceae, developing resources in understudied conifer families is essential for meaningful comparative genomic work that could further inform conservation and management for iconic species..

## ACKNOWLEDGEMENTS

This project was supported by a grant from Save The Redwoods League for the Redwood Genome Project (to DN), and by grants from the National Institute of Food and Agriculture of the U.S. Department of Agriculture (http://nifa.usda.gov; 2018-67011-28025 to AR and 2018-67015-28199 to AZ). Illumina and PacBio sequencing were carried out by the DNA Technologies and Expression Analysis Cores at the UC Davis Genome Center, supported by NIH Shared Instrumentation Grant 1S10OD010786-01. Part of this research project was conducted using computational resources at the Maryland Advanced Research Computing Center (MARCC) and at the Computational Biology Core, Institute for Systems Genomics, University of Connecticut. Professor Stephen C. Sillett and his group at Humboldt State University made this project possible by climbing SEGI 21 and obtaining cones and foliage for sequencing. Marc Crepeau’s skill at megagametophyte dissection, DNA extraction, and library prep is well appreciated. Bill Libby provided valuable support for this project, in the form of scientific guidance and both enthusiasm and expertise in giant sequoia genetics. Thank you to Sequoia/Kings Canyon National Park for allowing us to conduct research inside the park.

---

**Supplemental Figure 1:**
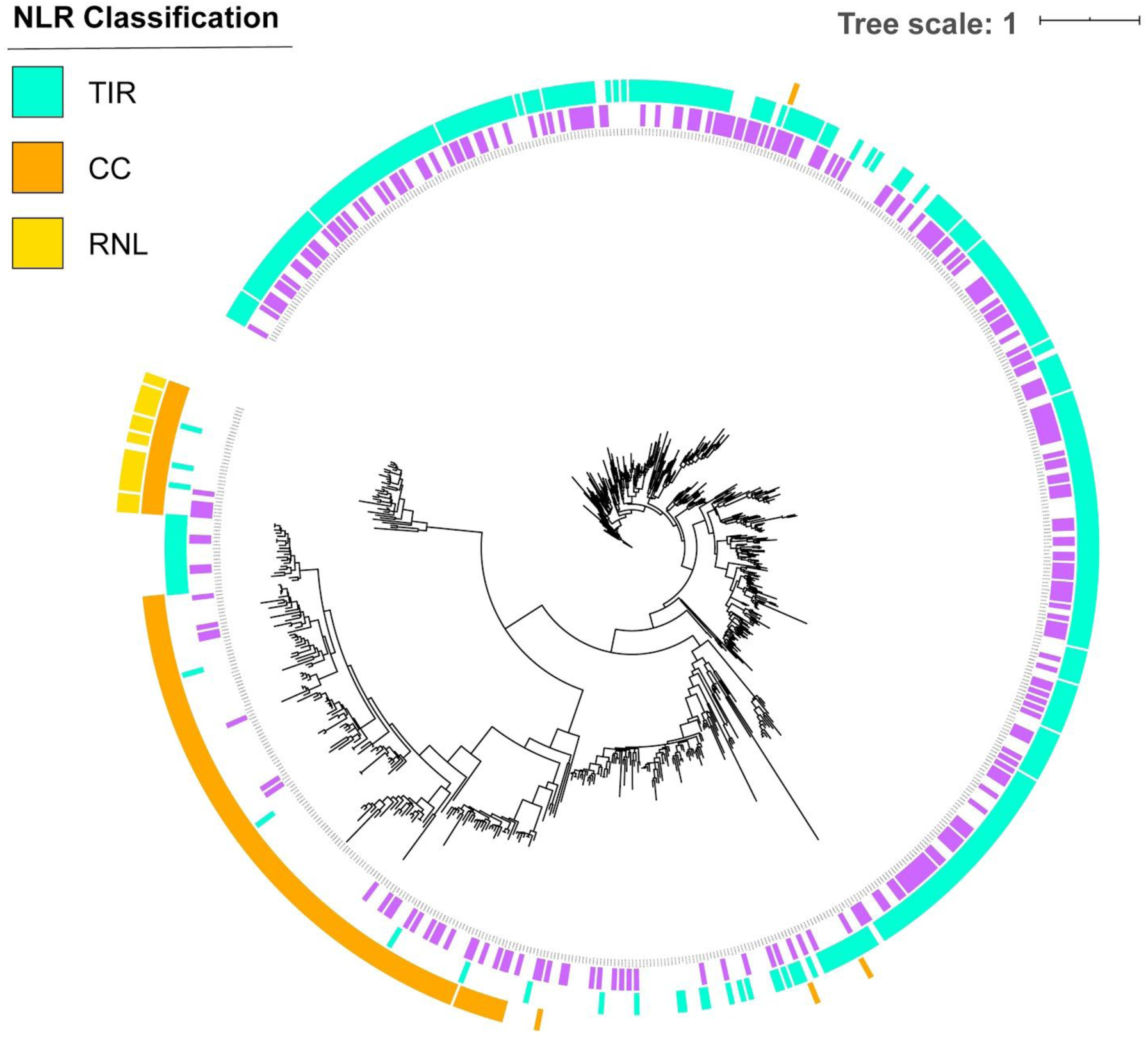
Maximum likelihood tree of NB-ARC domains of all NLR-Annotator detected NLR genes. The purple ring represents consensus NLR genes. N-terminal domains are indicated in the outer rings (TIR- light blue, CC- orange, RNL- yellow).

**Table S1.** Completeness of conifer genome assemblies assessed with BUSCOv3.0.2. Giant sequoia 2.0 is consistent with completeness of other conifer assemblies.

|  | *Sequoiadendron giganteum* | *Picea glauca* | *Picea abies* | *Pinus lambertiana* | *Pinus taeda* | *Pseudotsuga menziesii* |
| --- | --- | --- | --- | --- | --- | --- |
|  | Giant sequoia | White spruce | Norway spruce | Sugar pine | Loblolly pine | Doug fir |
| Complete BUSCOs (C) | 611 | 443 | 505 | 396 | 636 | 484 |
| Complete and single-copy BUSCOs (S) | 575 | 316 | 434 | 349 | 508 | 412 |
| Complete and duplicated BUSCOs (D) | 36 | 127 | 71 | 47 | 128 | 72 |
| Fragmented BUSCOs (F) | 192 | 182 | 150 | 172 | 102 | 110 |
| Missing BUSCOs (M) | 811 | 815 | 785 | 872 | 702 | 846 |
| Total BUSCO groups searched | 1614 | 1440 | 1440 | 1440 | 1440 | 1440 |
| Percentage found | 37.86% | 30.76% | 35.07% | 27.50% | 44.17% | 33.61% |

**Table S2.** Classification and associated percentage of genome masked by repetitive elements in giant sequoia 2.0

| **Repeat Class** | **Masked % of genome** |
| --- | --- |
| **DNA** |  |
| DNA/CMC-EnSpm | 0.36161 |
| DNA/MuLE-MuDR | 1.55370 |
| DNA/Sola | 0.77113 |
| DNA/TcMar-Fot1 | 0.05913 |
| DNA/hAT-Tag1 | 0.47514 |
| DNA/hAT-Tip100 | 0.13902 |
| *DNA total* | 3.35974 |
| **LINE** |  |
| LINE/L1 | 1.77165 |
| LINE/L1-Tx1 | 0.40261 |
| LINE/Penelope | 0.02434 |
| LINE/RTE-X | 0.11875 |
| LINE/Tad1? | 0.00674 |
| *LINE total* | 2.32409 |
| **LTR** |  |
| LTR | 0.21792 |
| LTR/Copia | 8.05066 |
| LTR/ERVK | 0.46223 |
| LTR/Gypsy | 19.62522 |
| *LTR total* | 28.35604 |
| Low_complexity | 0.16773 |
| RC/Helitron | 0.07451 |
| Satellite | 0.00005 |
| Simple_repeat | 2.03401 |
| Unknown | 42.34551 |

**Table S3.**
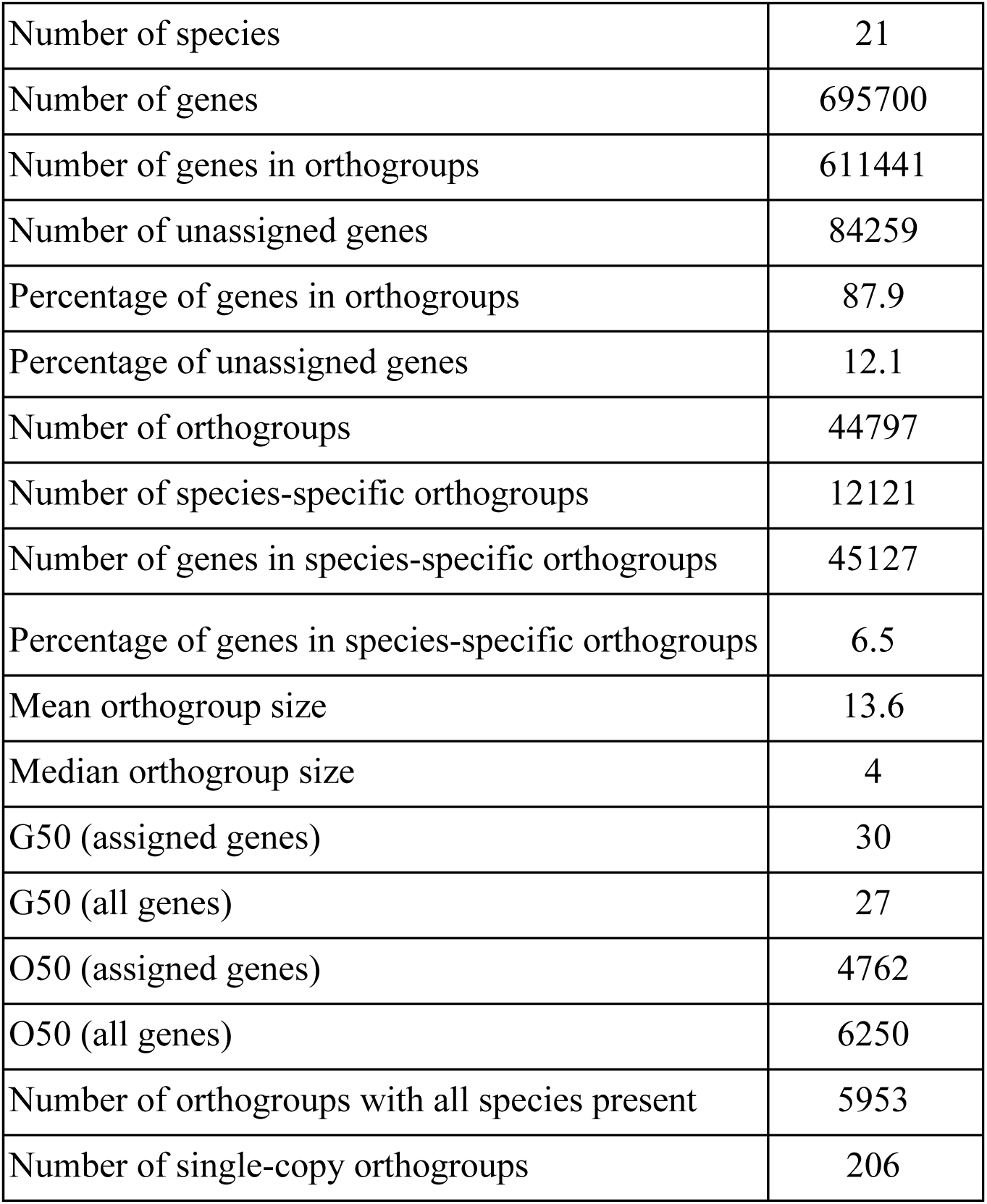
Orthogroup clustering of 695,700 protein sequences from twenty gymnosperms plus an outgroup (*Amborella trichopoda*)

**Table S4.**
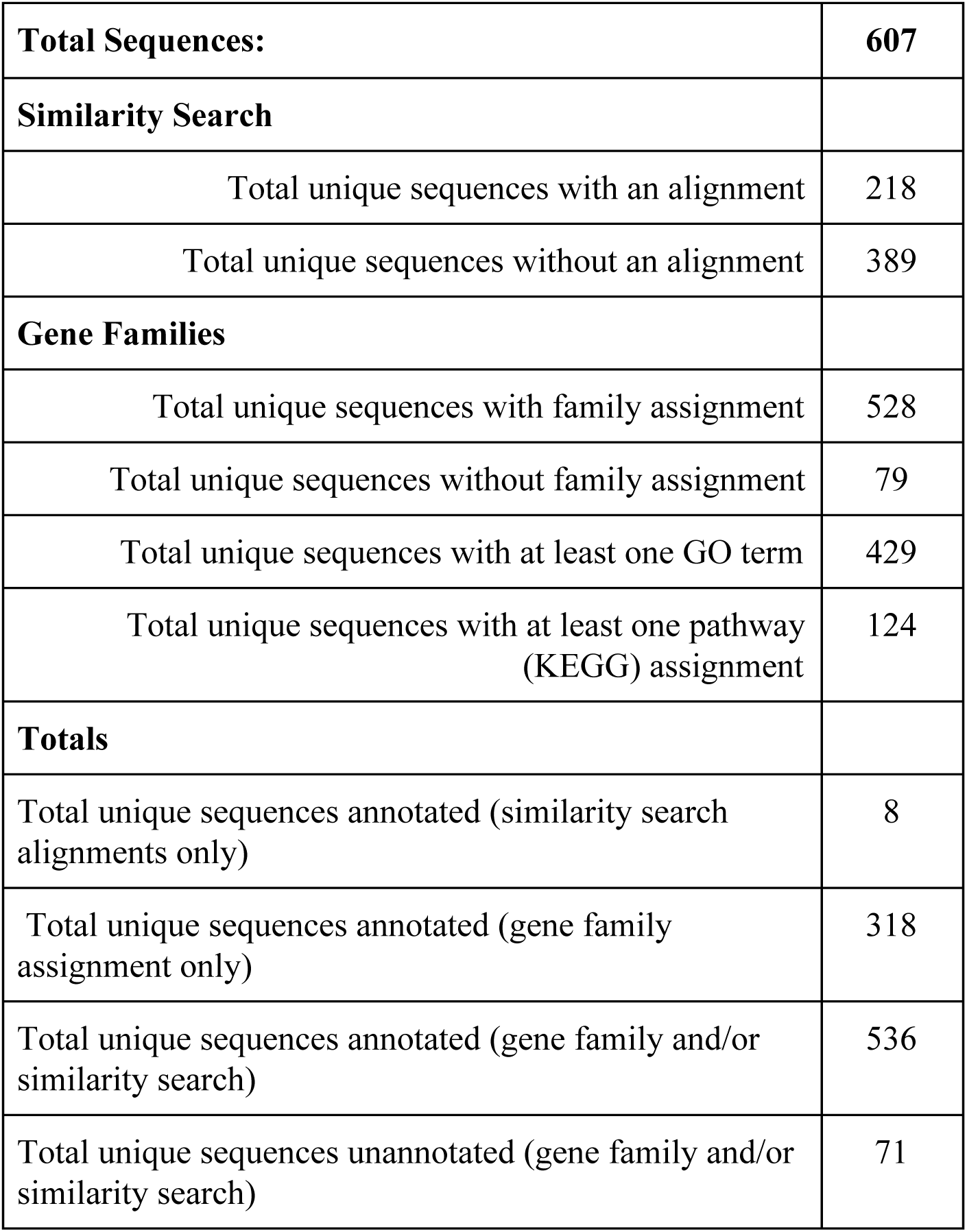
Annotation summary for 607 species-specific giant sequoia orthogroups

|  |  |
| --- | --- |
| **Total Sequences:** | **607** |
| **Similarity Search** |  |
| Total unique sequences with an alignment | 218 |
| Total unique sequences without an alignment | 389 |
| **Gene Families** |  |
| Total unique sequences with family assignment | 528 |
| Total unique sequences without family assignment | 79 |
| Total unique sequences with at least one GO term | 429 |
| Total unique sequences with at least one pathway<br>(KEGG) assignment | 124 |
| **Totals** |  |
| Total unique sequences annotated (similarity search<br>alignments only) | 8 |
| Total unique sequences annotated (gene family<br>assignment only) | 318 |
| Total unique sequences annotated (gene family and/or<br>similarity search) | 536 |
| Total unique sequences unannotated (gene family and/or<br>similarity search) | 71 |

